# Decoding cell identity with multi-scale explainable deep learning

**DOI:** 10.1101/2024.02.05.578922

**Authors:** Jun Zhu, Zeyang Zhang, Yujia Xiang, Beini Xie, Xinwen Dong, Linhai Xie, Peijie Zhou, Rongyan Yao, Xiaowen Wang, Yang Li, Fuchu He, Wenwu Zhu, Ziwei Zhang, Cheng Chang

**Affiliations:** Tsinghua-Peking Joint Center for Life Sciences, School of Life Sciences, Tsinghua University, Beijing, 100084, China; State Key Laboratory of Medical Proteomics, Beijing Proteome Research Center, National Center for Protein Sciences (Beijing), Beijing Institute of Lifeomics, Beijing 102206, China; Department of Computer Science and Technology, Tsinghua University, Beijing, 100084, China; Department of Basic Medical Sciences, School of Medicine, Tsinghua University, Beijing, 100084, China; Tsinghua-Berkeley Shenzhen Institute, Tsinghua University, Shenzhen, 518055, China; International Academy of Phronesis Medicine (Guang Dong), No. 96 Xindao Ring South Road, Guangzhou International Bio Island, Guangzhou 510000, China; Center for Machine Learning Research, Peking University, Beijing 100871, China; AI for Science Institute, Beijing, 100084, China

## Abstract

Cells are the fundamental structural and functional units of life. Studying the definition and composition of different cell types can help us understand the complex mechanisms underlying biological diversity and functionality. The increasing volume of extensive single-cell omics data makes it possible to provide detailed characterisations of cell types. Recently, there has been a rise in deep learning-based approaches that generate cell type labels solely through mapping query data to reference data. However, these approaches lack multi-scale descriptions and interpretations of identified cell types. Here, we propose Cell Decoder, a biological prior knowledge informed model to achieve multi-scale representation of cells. We implemented automated machine learning and post-hoc analysis techniques to decode cell identity. We have shown that Cell Decoder compares favourably to existing methods, offering multi-view interpretability for decoding cell identity and data integration. Furthermore, we have showcased its applicability in uncovering novel cell types and states in both human bone and mouse embryonic contexts, thereby revealing the multi-scale heterogeneity inherent in cell identities.

## Introduction

Cells are the basic structural and functional units of life^1^. The complex functions of different tissues and organs are rooted in cellular composition, and studying the organisation and function of various cells can help understand how organisms achieve normal life functions. Cell types have been defined according to their structures and functions for centuries. Usually, the same kind of cells exhibit similar characteristics and functions. Classifying and annotating cells can significantly aid in understanding their organisation and functions^2^. The increasing application of single-cell transcriptomic technologies in biological research has greatly advanced the study of cell types^3–6^. However, the varying properties exhibited in different cell types at multiple scales present substantial challenges for precise cell-type definition and annotation.

The identification of cell types in single-cell transcriptomics data is usually reliant on a multi-step process. It includes the preprocessing of transcriptomics profiles, dimensionality reduction, and unsupervised clustering. Subsequently, category annotation is conducted based on manually curated differentially expressed marker genes^7, 8^. Cell identification based on the traditional approach is time-consuming and laborious, and the selection of marker genes heavily relies on researchers’ domain knowledge, which is empirical and easily biased. As the accumulation of annotated single-cell transcriptomics data provides a large number of reference datasets for cell type identification, some representative deep learning models, including fully connected neural networks, autoencoders and transformers, are applied for mapping and migrating from reference datasets to new datasets for cell-type identification^9–12^. While the aforementioned methods have achieved commendable performances across different datasets, their ‘black box’ nature renders them largely unexplainable. The essence of the model learning process and human reasoning is significantly different, making it difficult to understand how deep learning models learn^13^ from single-cell data. However, for biological research, the transparency of the model is as important as its accuracy. A clear understanding of a model’s workings is indispensable for interpreting the biological significance of its findings.

A significant trend in machine learning is the development of explainable deep learning methods (XAI)^14^. For instance, the incorporation of domain knowledge into the model for drug response prediction, tumour typing and biomarker discovery^15–19^. However, these methods do not fully leverage biological domain knowledge, particularly the interactions between proteins and the interdependencies among biological pathways. Therefore, constructing a multi-scale interpretable model for cell-type identification remains a challenge.

Here, we propose an interpretable deep learning model called Cell Decoder, which embeds multi-scale biological knowledge into the graph neural network, enabling the decoding of distinct cell identity features. Cell Decoder constructs a hierarchical graph structure based on the interactions between genes, the mapping relationships between genes and pathways, and the hierarchical pathway information. Through the application of automated machine learning techniques, the model’s representation power is enhanced, facilitating precise and robust cell-type identification and multi-scale data integration. Moreover, we have developed a multi-view posterior probability interpretation method, elucidating the model’s learning and decision-making processes and mapping them to biological explanations. Cell Decoder facilitates the understanding of the interactions, pathways and biological processes that distinguish different cell types, providing significant implications for deeper exploration of cell identity and function.

## Results

### Overview of Cell Decoder

Traditional deep neural networks exhibit heightened capacity at the cost of reduced model interpretability. Here, we aim to design a model that can maintain robust representational power while offering biologically interpretable insights for identifying cell identity features. Cell Decoder designs an explainable graph neural network to model the multi-scale biological interactions and gene expressions for cell-type identification.

First, Cell Decoder leverages biological domain knowledge from curated databases and gene expressions as inputs. This biological domain knowledge includes protein-protein interaction (PPI) networks, gene-pathway maps and pathway-hierarchy relationships (Fig.1a). Cell Decoder processes these relationships to construct multi-scale interactions, including the gene-gene graph, gene-pathway graph, pathway-pathway graph, pathway-biology process (BP) graph and BP-BP graph (Fig.1b), represented as graph structures and fed into the model as inputs. Concurrently, gene expressions are used as features for each node.

To integrate information within the same resolution or across different scales effectively, Cell Decoder designs intra-scale and inter-scale message passing layers respectively (Fig.1b). The former shares messages within homogeneous biological entities such as different genes, pathways or BPs while the latter aggregates information from a fine-grained resolution to a coarse-grained one, i.e. from genes to pathways or from pathways to BPs. Then, Cell Decoder utilises mean pooling to summarise the node representations of the BPs in the last graph layer into cell representations and adopts a multi-layer perceptron classifier for cell-type identification. Cell Decoder is trained end-to-end by minimising the cross-entropy loss between predicted and ground-truth cell labels.

To adapt to various intricate cell-type identification scenarios, Cell Decoder utilises an automated machine learning (AutoML) module to search the model design automatically, encompassing the choices of intra-scale and inter-scale layers, hyper-parameters and architecture modifications. The searched Cell Decoder instantiation is specifically tailored to fit the targeted cell-type identification scenario, consequently leading to improved results.

Lastly, to provide model interpretability and gain insights into the identified cell types, Cell Decoder incorporates post-hoc interpretability modules (Fig.1c). Through hierarchical Gradient-weighted Class Activation Mapping (Grad-CAM)^20^ analysis of the fitted model, a diverse set of biological features can be identified, including pathways and biological processes crucial for predicting different cell types. This provides a multi-view biological characterisation that enhances our ability to decode cell identity. Moreover, leveraging attention scores can further differentiate cell types based on the PPI network within cells (Methods).

### Cell Decoder achieves superior performances and robustness for cell-type identification

We benchmarked Cell Decoder using 7 different datasets, including human blood (HU_Blood)^21^, human bone marrow (HU_Bone)^6^, human liver (HU_Liver)^22, 23^, human kidney (HU_Kidney)^24^, human pancreas (HU_Pancreas)^25–27^, mouse lung (MU_Lung) and mouse pancreas (MU_Pancreas)^28^ (Supplementary Data 1) against 9 popular cell identification methods^9, 11, 12, 29–34^ on its prediction accuracy and Macro F1 score. Macro F1 score here is defined as the Macro average of the F1 scores for each cell type. Considering the prevalence of a few cell types within the entirety of a single-cell dataset, the Macro F1 score is more suitable than mere accuracy for evaluating the model’s capability in recognising various cell types.

Cell Decoder ranked first for the average of both Macro F1 score (Fig. 2a) and accuracy across all datasets (Supplementary Fig. 1a and Supplementary Data 2). Compared to the second-best deep learning method (ACTINN Macro F1 score at 0.72), Cell Decoder showed a 12.5% improvement in Macro F1 score. In each dataset, Cell Decoder has achieved the best performance in terms of the Macro F1 metric. Considering the inherent noise in single-cell datasets, the feasibility of model transfer has been limited. To gauge the capacity of fitted model transferring across diverse datasets, we conducted feature perturbation experiments (Methods). These experiments introduced random noise with varying rates of perturbations into the test data. As the level of data perturbation increases, all models exhibit a certain degree of decline in prediction performance. Compared to other models with transfer capabilities, Cell Decoder has demonstrated remarkable improvements in robustness across all 7 datasets (Fig. 2b). This indicates that Cell Decoder can recognise the efficient identity of different cell types and has strong resistance to data noise.

For cell identification, dealing with imbalanced cell-type proportions within datasets and distribution shifts between reference and query datasets are common challenges. Therefore, evaluating the performance of models under these two scenarios is significantly important. Imbalanced distribution of epithelial cell types in the MU_Lung. AT2 cells make up 82% of the reference data, while AT1 cells, Ciliated cells, and Club cells account for 8%, 8%, and 2%, respectively (Fig. 2c). Cell Decoder outperforms other deep learning models in predicting accuracy for the four imbalanced cell types (Fig. 2d). In scenarios with imbalanced cell types of Endothelial cells, Immune cells and Mesenchymal cells in MU_Lung, Cell Decoder also achieves the highest prediction accuracy for the minority cell types (Supplementary Fig. 1b-g). In the HU_Liver dataset, there is a clear data shift, with the proportions of cell types in the reference and query datasets exhibiting opposite trends (Fig. 2e). Hepatocyte, Plasma, Mono/Macro and Portal endothelial (highlighted in red in Fig. 2f) have a higher proportion in the query dataset compared to the reference dataset. Cell Decoder achieves a recall of 0.88 on the query dataset, marking a 14.3% improvement over the second-best method, ACTINN, which achieves a recall of 0.77. In comparison, Cell BLAST and TOSICA have recalls of 0.69 and 0.68, respectively. We also evaluated the performance of the deep learning models in terms of precision and Macro F1 scores. Cell Decoder achieves a precision of 0.86 and a Macro F1 score of 0.85, demonstrating significant improvements of 11.7% and 23.2% over the second-best model (Supplementary Fig. 1h-i).

We then conducted ablation experiments on the model by randomly removing biological prior knowledge (nodes and edges in the graph) and testing the retrained model (Methods). When the perturbation rate is 100%, it implies that the edges of the model are fully removed, with the exception of self-loop. As the rate of graph node perturbation increases, model performance decreases in almost all datasets, except HU_Pancreas and MU_Lung, where it remains relatively stable (Supplementary Fig. 2a). When randomly perturbing the edges of the model, there is also a noticeable decrease in prediction performance. In the case of the MU_Lung dataset, the Macro F1 score and accuracy slightly decrease. For both HU_Blood and HU_Bone, the Macro F1 score and accuracy initially decline but eventually exhibit a modest recovery (Supplementary Fig. 2b). This suggests that biologically informed modelling improves deep learning performance of predicting cell type.

### Cell Decoder enables multi-scale data integration without batch labels

In-depth cell identification requires integration across multiple datasets. However, batch effects can arise in different datasets due to factors such as varying experimental protocols. Cell Decoder can effectively identify cell types, enabling the integration of diverse datasets without the requirement of batch labels. By incorporating multi-scale biological prior knowledge into Cell Decoder (Fig. 1a), we can obtain low-dimensional embedding of cells at pathway and biological process layers. We evaluated the multi-scale data integration capability of Cell Decoder on the human immune cell dataset^35–39^ with 10 different batches provided by scIB^40^ in comparison with Harmony^41^, Scanorama (embedding)^42^, and scVI^43^. The data integration performance of Cell Decoder is illustrated by the pathway embedding and biological process (BP) embedding (Fig. 3a, b and Supplementary Fig. 3). Cell Decoder effectively removed batch effects between individuals and platforms, while preserving biological variations. We evaluated the performance of these methods using a total of eight metrics, divided into two categories: batch correction and biological variation preservation. The embedding at pathway layer, i.e., Cell Decoder (Pathway) achieved the best results on batch ASW, kBET and graph iLISI metrics, indicating its superior batch correction performance. Moreover, the embedding at BP layer, i.e., Cell Decoder (BP) demonstrated significant improvements over other methods in preserving biological variation at NMI cluster, ARI cluster, cell type ASW and isolated label F1 metrics (Supplementary Data 3). On an overall basis (the average of all metrics), Cell Decoder (BP) achieved a 20.5% improvement (Fig. 3c) compared with existing methods.

**Fig. 1.**
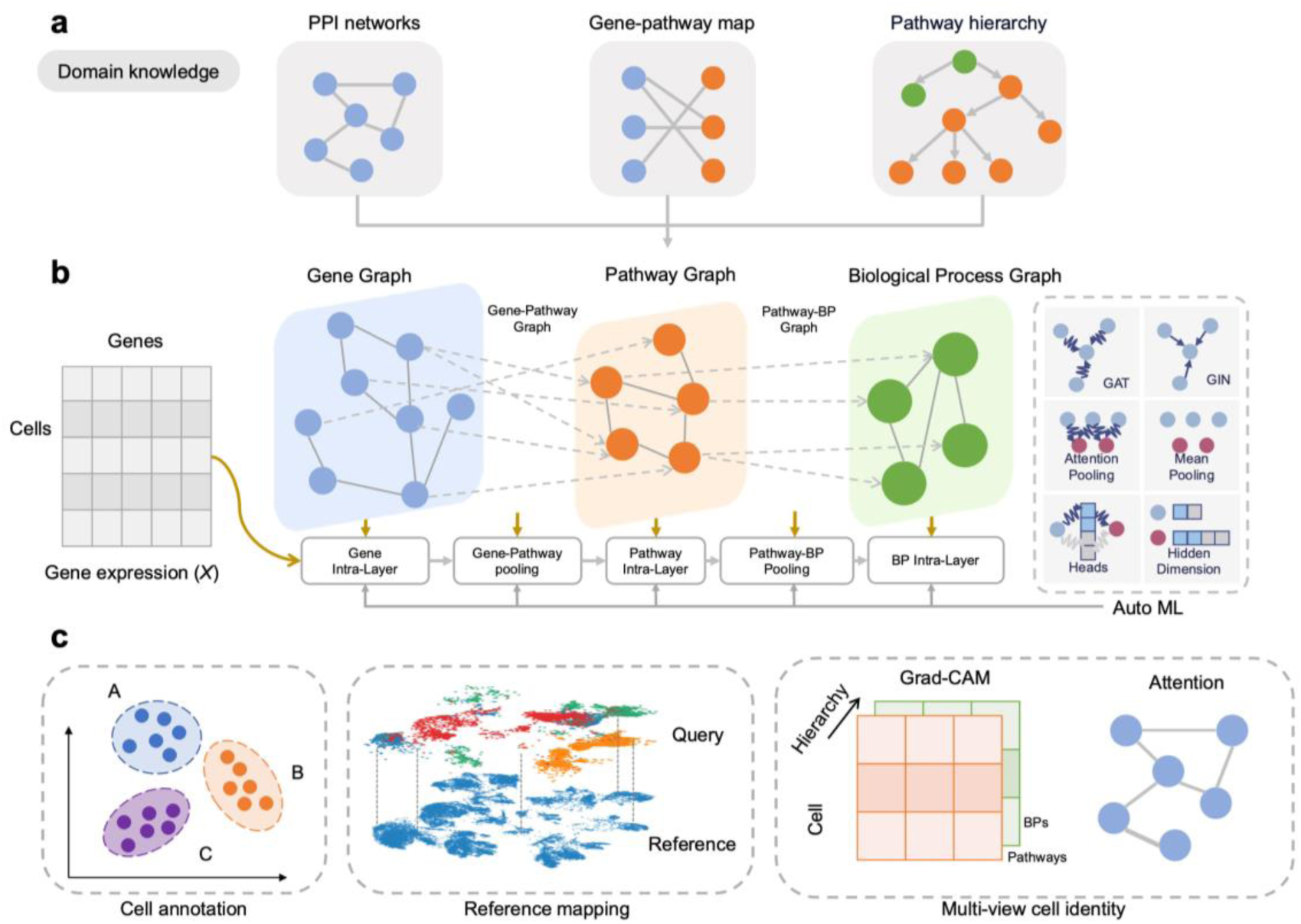
An overview of Cell Decoder. a. Cell Decoder explicitly leverages biological domain knowledge in cell-type identification, which consists of protein-protein-interaction networks, gene-pathway maps, and pathway-hierarchy relationships. These networks are represented as graphs (adjacency matrices) that serve as inputs to Cell Decoder. b. In addition, Cell Decoder incorporates gene expressions as features for each cell. Cell Decoder employs intra-scale and inter-scale messaging passing layers to integrate information across different scales, thereby obtaining the representations of genes, pathways, and biological processes. The specific layers, as well as other hyper-parameters and architecture modifications, are automatically designed using AutoML techniques to adapt to various cell-type identification scenarios. GAT: Graph Attention Networks; GIN: Graph Isomorphism Networks. c. The output of Cell Decoder can be applied for cell annotation, reference mapping and incorporates post-hoc interpretability modules to provide explainable multi-view cell identity. Grad-CAM: Gradient-weighted Class Activation Mapping.

**Fig. 2.**
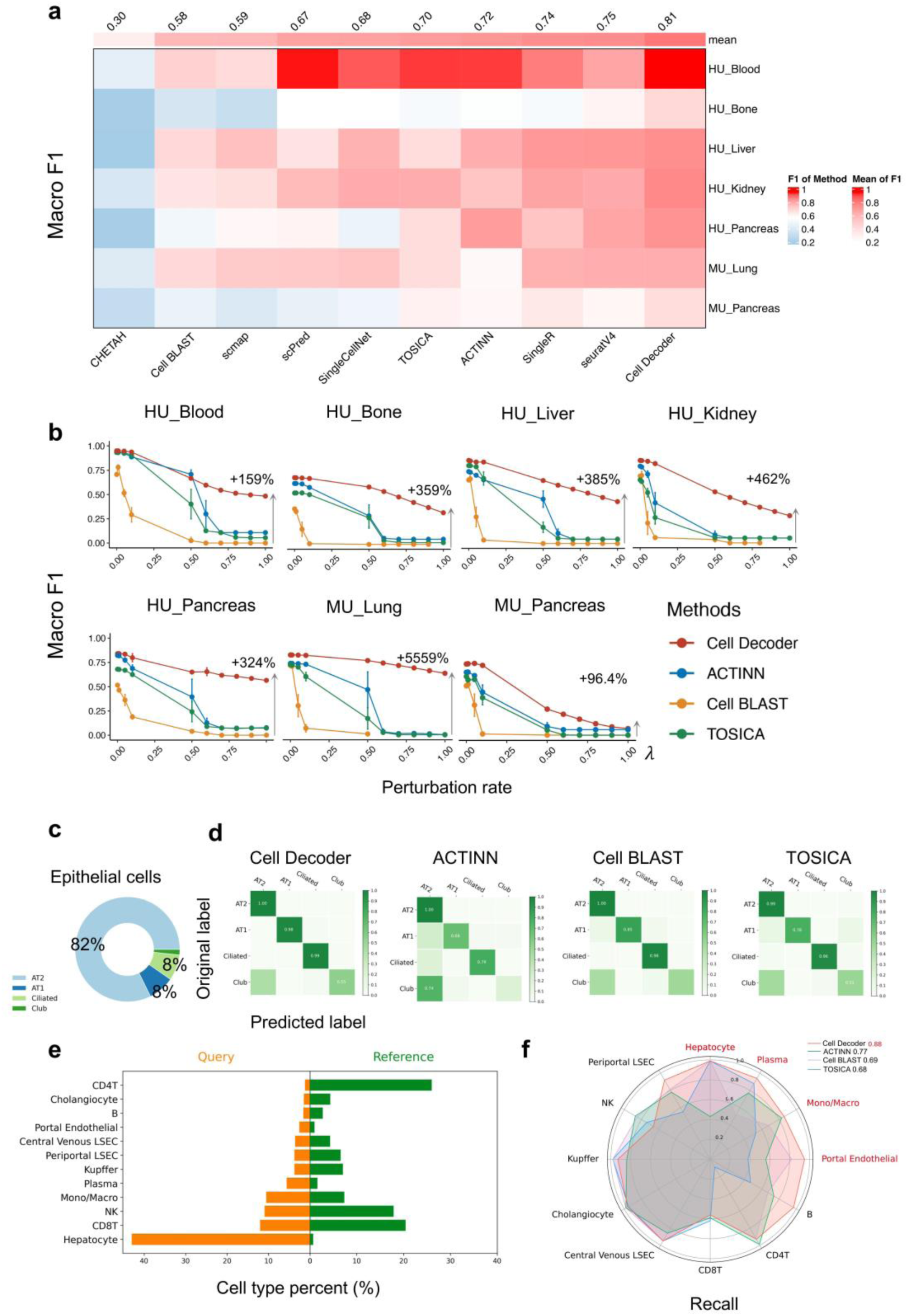
Systematic benchmarks and robustness evaluation of Cell Decoder for cell identification. **a** Macro F1 score for different cell identification methods on human and mouse datasets. Columns are sorted by the mean Macro F1 of each method on all datasets (top). **b** Robustness evaluation for cell identification based deep learning methods at 10 different feature perturbation rates (0, 0.01, 0.05, 0.1, 0.5, 0.6, 0.7, 0.8, 0.9, and 1). The average improvement ratio for Cell Decoder compared to the second-best method was shown in each dataset. (*n* = 3 repeats with different feature noise random seeds, *λ*: the weight of injected noise. See Methods). **c** Imbalanced epithelial cell type ratio in mouse lung. **d** Confusion matrix of different cell identification methods for predicting imbalanced epithelial cell type (**c**). Values are normalised within each row, and values greater than 0.5 are noted. **e** Data shifted between reference and query in human liver dataset. **f** Recall score of each cell type (indicated in red if its proportion in the query is greater than that in the reference), and the average recall scores for each method are labelled on the upper right corner.

**Fig. 3.**
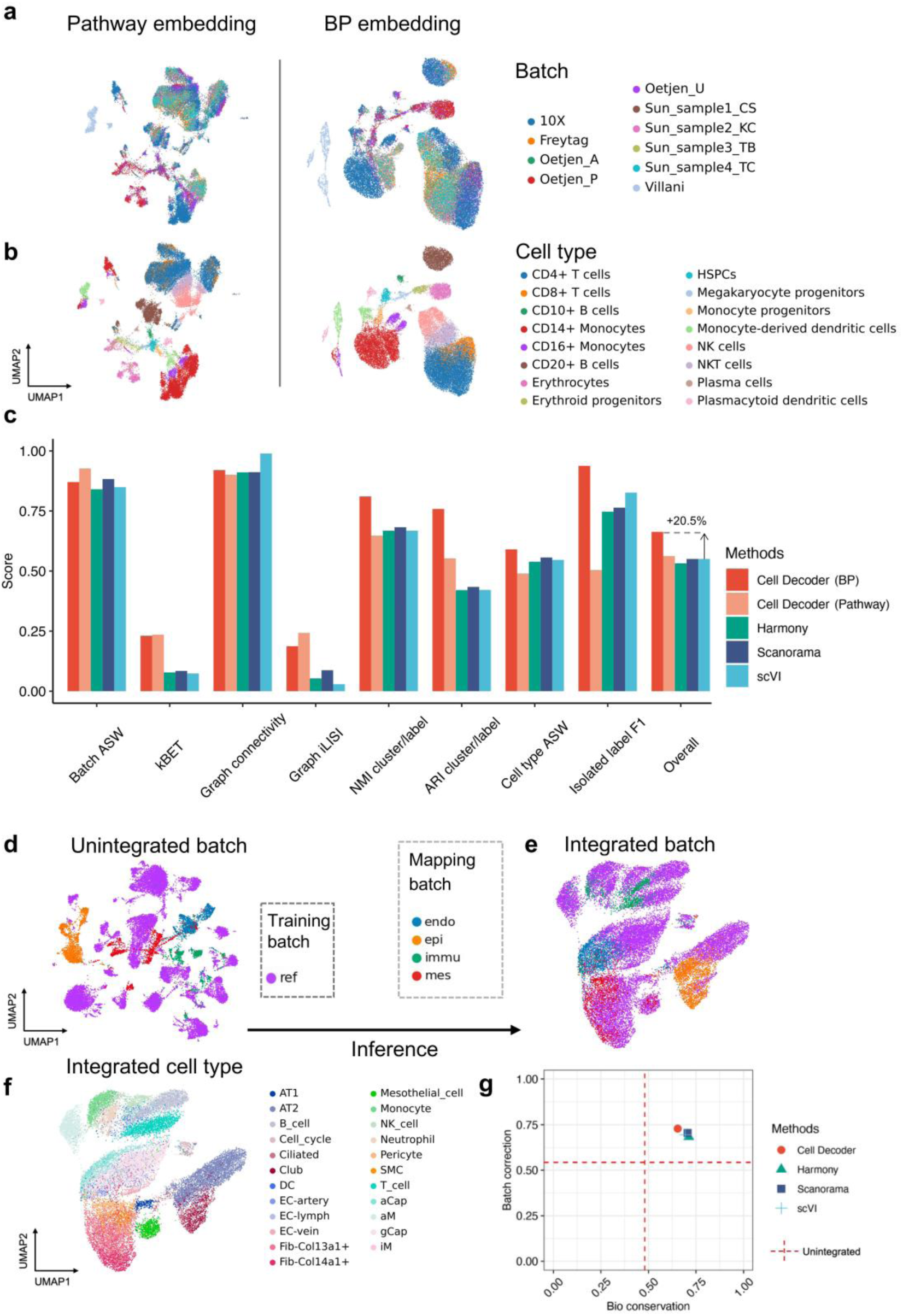
Multi-scale biologically informed data integration. **a, b** UMAP visualisation of the multi-scale integrated cell embeddings for pathway (left) and biological process (right) by cell identity (**a**) and batch annotation (**b**). **c** Comparison of data integration accuracy across different models without batch labels. The metrics measure bio-conservation and batch correction. Overall scores (the last line) are computed on the basis of the mean of all metrics. **d-g** Pretrain on reference batch then integrate query batches using the fitted model. UMAP visualisation of unintegrated batch (**d**), integrated batch (**e**) and integrated cell type (**f**). Scatter plot of the mean overall batch correction against mean overall bio-conservation score for the different methods, and the red dotted lines indicated unintegrated score (**g**).

Similar to domain generalization techniques^44^, Cell Decoder can generalize to a new batch that it has never seen during training. Namely, with a pre-trained model, all data can be embedded into the same low-dimensional space without any finetune process, which significantly enhances the efficiency and the usability of the tool for further application. Furthermore, Cell Decoder can extract generalizable representations from a single batch. The MU_Lung dataset comprises five batches (Reference, Endothelial cells, Epithelial cells, Immune cells, and Mesenchymal cells), among which Reference (ref) was used as the training set, while Endothelial cells (endo), Epithelial cells (epi), Immune cells (immu), and Mesenchymal cells (mes) were employed as the test sets. The UMAP visualisation highlighted the batch effects in the MU_Lung raw data. Meanwhile, the integration results in a low-dimensional space showed the ability of Cell Decoder to remove these batch effects (Fig. 3d-f). Compared to the other three data integration methods, Cell Decoder, pre-trained on only one batch of data, outperformed in batch effect correction but exhibits slightly weaker performance in biological variation preservation (Fig. 3g). However, Cell Decoder retains the unique capability of providing cell-type identification results, which sets it apart from the other methods.

### Discovery of novel cell types and cell states using Cell Decoder

One significant challenge in cell identification tasks is discovering cell types within the query dataset that are absent from the reference dataset. The majority of existing methods tend to categorise novel classes by forcefully aligning them with the closest known class. Such approaches are not able to discover novel cell types or cell states in the query dataset. Benefiting from biologically informed modelling, Cell Decoder possesses significant potential for capturing subtle differences between novel and known cell types. By predicting the probabilities of different cell types, it can automatically uncover potential new cell types. Moreover, it can decode the identity features of new cell types based on prior knowledge and post-hoc analysis (Methods).

To verify Cell Decoder’s ability to discover novel cell types, we masked ‘Mono/Macro cells’ in the training dataset HU_Kidney, while this cell type exists in the test set, thus simulating the scenario of encountering new cell types in the query dataset (Fig. 4a). Cell Decoder provides the predicted probabilities for each cell. If the highest probability falls below a threshold (0.95), the cell is classified as ‘Novel’, suggesting it is likely a new cell type not included in the training set (Fig. 4a right). Despite their high predictive accuracies, the methods such as Seurat, SingleR, and ACTINN (Fig. 2a) are unable to automatically identify newly emerged cell types in the query dataset. Instead, they forcibly categorise them as existing cell types in the reference. TOSICA (cutoff=0.95) and Cell BLAST (P<=0.05) also identify novel cells in the query dataset by predicting the probabilities of different cells. On the masked Mono/Macro cell type, Cell BLAST achieves a recall of 0.20, while the remaining Mono/Macro cells are predicted as CD4T, CD8T cells, with a small portion being labeled as ambiguous (Fig. 4b middle). TOSICA correctly labels 37% of Mono/Macro cells but predicts a larger portion of cells as B cells (Fig. 4b right). In contrast, Cell Decoder achieves a recall of 0.94 for Mono/Macro cells, correctly identifying the vast majority of Mono/Macro cells in the query dataset. This represents a significant improvement compared to the other two methods (Fig. 4b left).

**Fig. 4.**
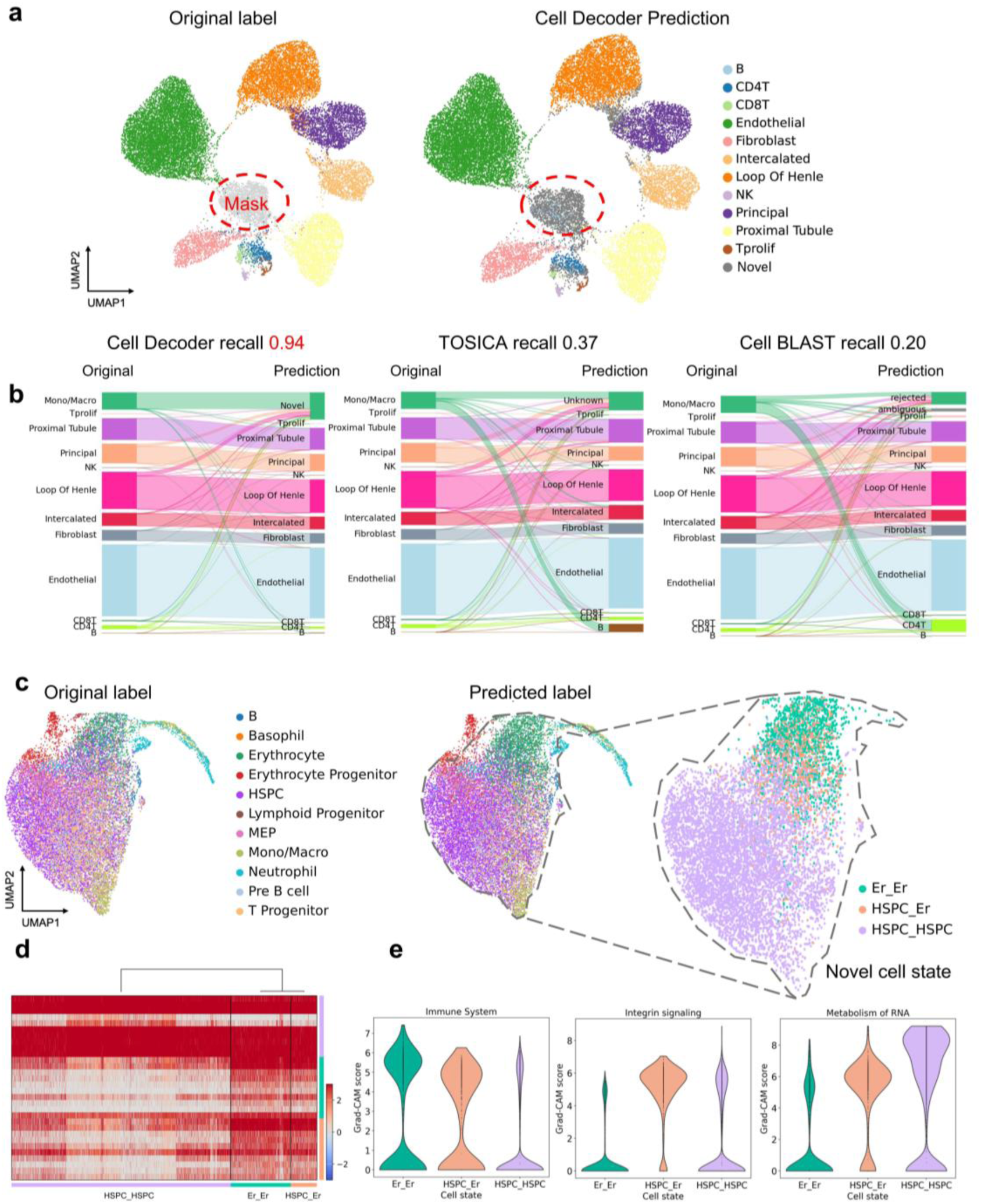
Identification of novel cell states and cell type. **a, b** Performance of different methods on the human liver for predicting novel cell type by masking Mono/Macro cell during model training (**a**). The recall of prediction novel cell type (**b**). Sankey plots comparing predictions on known and novel cell types across different methods (Left: original labels. Right: predicted labels). **c-e** Identification of novel cell state in the human bone dataset. UMAP visualisation of some labelled HSPC cells which are predicted as erythrocyte by Cell Decoder (Re-label as HSPC_Er in the left). Er_Er label indicates that both the original cell label and the predicted label are erythrocyte and the same applies to HSPC_HSPC (**c**). DEGs heatmap of HSPC_HSPC, HSPC_Er and Er_Er is hierarchically clustered (**d**). Violin Plots show the Grad-CAM score of three representative biology processes (immune system, integrin signaling and metabolism of RNA) in different cell states (**e**).

Identifying different cell states is also crucial, as a cell type may exist in various states^45^. Transcriptomic changes during cell state transitions are often more continuous than cell types. Consequently, in low-dimensional embedding spaces, these cells tend not to form distinct clusters but rather exhibit a continuous distribution. Manual identification methods relying solely on marker genes often face the challenges in identifying different cell states within the same cell type. However, Cell Decoder, as an automated cell identification method, is not dependent on specific marker genes. Instead, it is built upon biological prior knowledge, facilitating the extraction of cell identity features. This presents a promising potential for uncovering various cell states. In the HU_Bone dataset, some multipotent hematopoietic stem and progenitor cells (HSPC) cells were predicted as Erythrocytes by Cell Decoder (Fig. 4c). We re-labelled this cell type as HSPC_Er. Cells predicted to be consistent with the original labels of HSPC and Erythrocytes were marked as HSPC_HSPC and Er_Er, respectively. HSPC_HSP, HSPC_Er, and Er_Er exhibited a continuous change trend in the UMAP, with HSPC_Er positioned in an intermediate state between the two (Supplementary Fig. 4). We calculated the differential genes for the three cell types and performed hierarchical clustering (Fig. 4d). The differentially expressed genes showed that HSPC_Er cells were more similar to Er_Er cells and Er_Er cells exhibited the highest activation level in immunological processes, whereas HSPC_HSPC cells showed a more pronounced activation in the metabolism of RNA process. HSPC_Er cells showed the highest activation score in the integrin signalling pathway (Fig. 4e).

### Cell Decoder reveals cellular dynamic changes in mouse embryogenesis

Understanding the lineage relationships between cells and cell types, as well as the molecular programs governing the emergence of each cell type, constitutes a fundamental goal in developmental biology. For the mouse embryogenesis dataset^46–49^, the entire embryo has been profiled using scRNA-seq. However, systematically integrating the associated data remains a challenge. This challenge is attributed to technical issues such as varying technologies and batch effects, as well as the complexity of mouse development^50^. We integrated data from three stages of mouse embryo development at E6.25, E6.5, and E6.75, obtained from different technologies (Supplementary Data 4). In the mouse embryogenesis dataset, there are a total of seven cell types, namely epiblast (3302 cells), extraembryonic ectoderm (1220 cells), primitive streak and adjacent ectoderm (1214 cells), extraembryonic visceral endoderm (606 cells), embryonic visceral endoderm (295 cells), nascent mesoderm (159 cells) and parietal endoderm (44 cells).

Cell Decoder successfully integrated data from different developmental stages and demonstrated its batch correction capability through UMAP visualisation computed on the biological process embeddings (Fig. 5a). We randomly partitioned the data, allocating 80% to the training set and 20% to the validation set for model fitting. Furthermore, in the UMAP plot, we annotated the cell-type predictions both by Cell Decoder and the original labels from the data (Fig. 5b). The visceral endoderm encompasses the extraembryonic visceral endoderm (ExVE) and the embryonic visceral endoderm (EmVE). We learned the differences between ExVE and EmVE (E6.5 and E6.75) across various biological processes using Cell Decoder. The notably higher Grad-CAM scores of EmVE in cell-cell communication and extracellular matrix categories, as opposed to ExVE, suggested a heightened activity in these biological processes. Furthermore, EmVE at the E6.75 stage exhibited a significant level of programmed cell death, while ExVE showed pronounced activity in the immune system (Fig. 5c). Cell Decoder learns representations of EmVE and ExVE in the BP embedding, enabling further clustering into four subtypes (Fig. 5d). At E6.5, C0 is the predominant subtype, followed by C1 and C3, while C2 is the smallest subtype. However, at E6.75, the highest proportion is observed for C1, followed by C2 cells (Fig. 5e). The different subtypes showed distinct differences in the expression of marker genes in EmVE and ExVE. *Cer1* and *Lefty1* are highly expressed in C3 and C2, while *Nodal* exhibits the highest expression in C3, and Fgf8 exhibits the highest expression in C2 (Fig. 5f). Cell Decoder elucidates the different activation levels of the four cell subtypes across various biological pathways using Grad-CAM scores. For example, C2 showed a high activation in the NODAL, MTOR, WNT and MAPK signalling pathways (Fig. 5g). Cell Decoder also provides interaction analysis based on attention mechanisms, revealing that the interactions related to anterior-posterior axis formation. For example, *Lefty1-Nodal* and *Fgf8-Otx2* exhibited the highest scores in the C2 subtype (Fig. 5h). Cell Decoder reveals the dynamic changes of different cells during mouse embryonic development and identifies distinct subtypes of cells with diverse developmental functions.

**Fig. 5.**
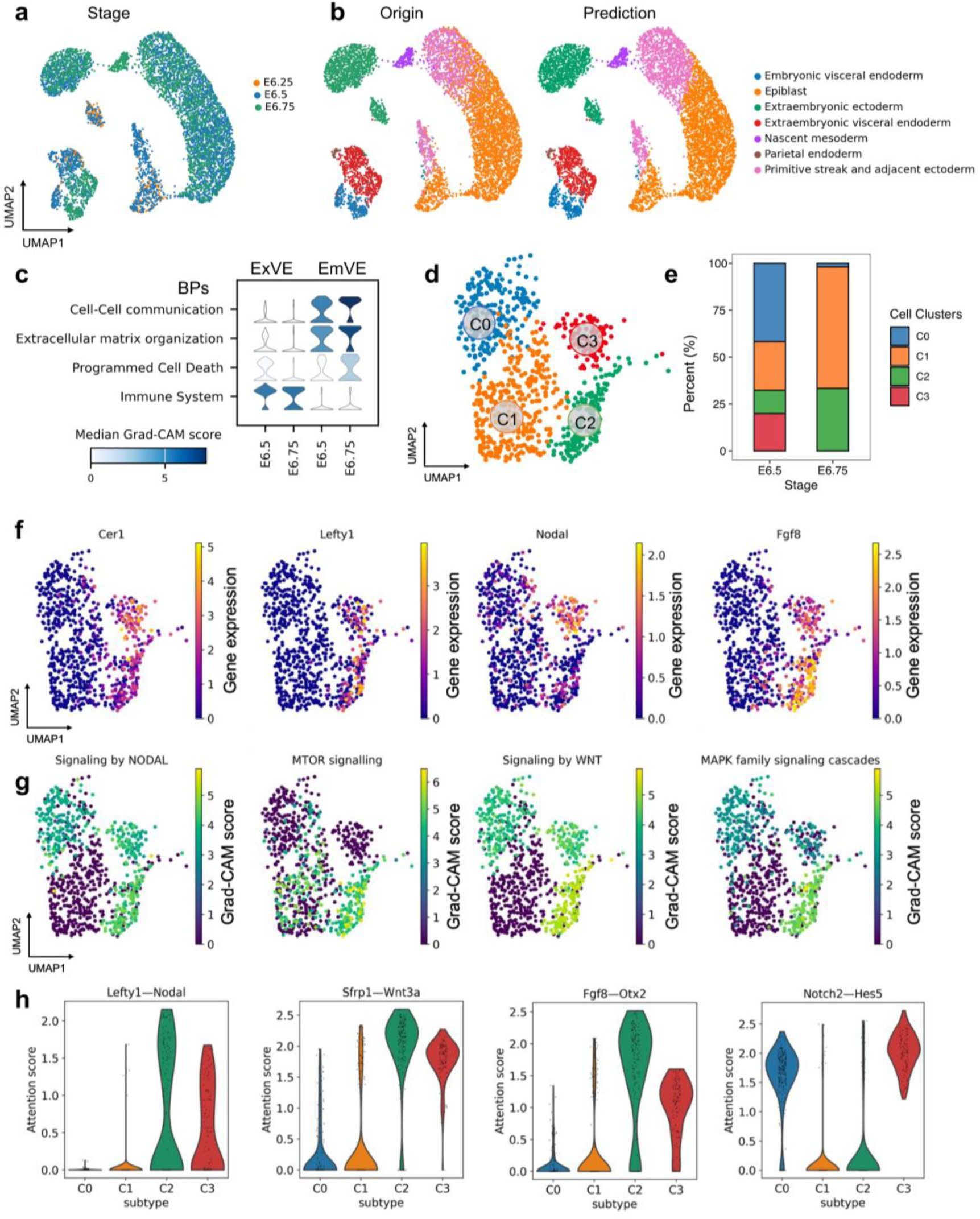
Decoding cellular dynamic change of mouse embryogenesis using Cell Decoder. **a, b** UMAP visualisation of the mouse embryo dataset based on Cell Decoder embedding coloured by stage (**a**) and cell type (**b**). **c** The activity of the biology processes across different stages for ExVE and EmVE based on Grad-CAM score. **d, e** Subtypes clustering of ExVE and EmVE based on Cell Decoder embedding (**d**) and its proportion of different stages (**e**). **f** Visualisation different marker genes distribution in ExVE and EmVE (**d**) of mouse embryo dataset. **g** UMAP representation subtypes of ExVE and EmVE (**d**) coloured by signaling pathways Grad-CAM score. **h** Violin Plots visualisation subtypes (**d**) for different interaction scores.

### Multi-view interpretability in Cell Decoder for cell identification

Due to the diverse properties exhibited by different cell types at multiple scales, defining, categorising, and understanding them pose significant challenges^45^. Probabilistic statistical and traditional deep learning-based cell identification methods can classify different cell types. However, most of these methods^9–12, 31^ lack interpretability. While TOSICA^9^ explains the mapping from gene to pathway through an attention mechanism, it falls short in decoding the identity features of a cell type. A cell type is generally considered to express an assemblage of cellular modules (protein complexes, pathways, and molecular machines constitute the structure and function of the cell) responsible for discrete subfunctions^51^. Cell Decoder leverages biological prior knowledge (protein-protein interactions, gene-pathways, biological process) to enhance the transparency of its network structure (Fig. 1a). Additionally, we have developed a multi-view post-hoc analysis (Methods) method to uncover the decision-making process of the model, thereby decoding different cell-type identities. In HU_Bone, Cell Decoder is capable of identifying differential biological process activations among different cell states (Fig. 4e). For HSPC, autophagy is crucial for maintaining their self-renewal capacity. Conversely, Erythrocytes play a role in immunity, acting as immune sentinels^52^. Beyond the hierarchical interpretation of gene-pathway-biological process relationships, Cell Decoder is also capable of explaining differences in protein-protein interaction pairs across different cell types through an attention mechanism (Fig. 1c). During mouse embryonic development, Cell Decoder learns a critical interaction pair (Lefty1-Nodal), revealing multi-level biological differences in the ExVM and EmVE cells (Fig. 5h). Cell Decoder provides a more comprehensive and in-depth perspective for defining and understanding different cell types or states.

## Discussion

We propose a novel biologically informed graph neural network architecture that integrates protein-protein interactions, gene-pathway mappings and hierarchical pathway information into a hierarchical graph neural network for decoding cell identity. This approach aims to simulate the intracellular gene/protein interactions and aggregate information from genes to pathways, then to biological processes. We utilised automated machine learning techniques to enhance model performance and streamline the cumbersome process of deep learning parameter tuning. Cell Decoder exhibits strong transparency and interpretability. Through multi-view posterior probability analysis of the fitted model, we gained deeper insights into different cell types across scales and interactions.

Cell Decoder has outperformed existing advanced methods across diverse benchmarks. In human and mouse cell identification tasks, Cell Decoder achieves accurate and stable knowledge transfer, demonstrating strong resilience against data noise and data shift. By learning cell-type-specific features, Cell Decoder achieves accurate data integration at different scales (pathways embedding and biological process embedding) without batch labels. We also explored a novel data integration approach, i.e., pretraining on one batch and transferring the model to others. This method not only accomplishes cell identification but also enables cross-batch data integration. Biologically informed modelling has effectively boosted the ability to discover new cell types or states.

Despite having the above advantages in decoding cell identity, Cell Decoder may face potential limitations in selecting biological knowledge for modelling. Our model leverages domain knowledge from multiple databases^53–55^ to enhance the interpretability of deep learning. However, the effectiveness of prior knowledge varies across different datasets. The results of knowledge ablation experiments on 7 datasets indicated that different tissues and organs required varying biological knowledge, demonstrating that the choice of biological knowledge modelling has a certain impact on the model’s performance. Another limitation involves the interpretation and validation of newly learned cell type identity features. While Cell Decoder captures novel changes in the query data, the validation of these changes at different scales requires additional domain expertise. Additionally, Cell Decoder relies on supervised training with labelled data, and its performance is constrained by the availability and quality of limited annotated data.

Meanwhile, with the development of single-cell multi-omics^56^ technologies that can capture protein expression levels within individual cells, Cell Decoder can naturally extend to multimodal datasets, thereby gaining more insights into cell-type regulation. Due to the incorporation of gene/protein interaction information, Cell Decoder opens up the possibility of predicting cell responses to genetic perturbations^57^. We believe this can bring new interpretability and biological insights to define and understand the identity features of different cell types.

## Methods

### Biologically informed matrix

We integrated STRING V11.5 (https://string-db.org), MSigDB (https://www.gsea-msigdb.org/gsea/msigdb) and Reactome (https://reactome.org) database to build the biologically informed hierarchy graph. We filtered the protein-protein interactions with high confidence (combinations score>=850) from STRING to construct the graph *G* = (*V*, *E*) where *V* is the set of protein nodes in graph *G*, and E is the set of edges between any two nodes in graph *G*. The adjacency matrix *A* is in the form a binary matrix with *A*_*i*,*j*_ = 1 if the i-th protein interact with jth protein and 0 otherwise. We use depth-first search (DFS) to covert the pathway-directed acyclic graph (DAG) to a scalable hierarchy network with parameter n layers. Then, we map the graph *G* to hierarchy pathways using Reactome and MsigDB. The mask matrix *M* with columns corresponding to pathways and rows corresponding to genes, with *M*_*i*,*j*_ = 1 if the ith gene is in the *j*_*t*ℎ_ pathway and 0 otherwise.

### The architecture of Cell Decoder

Cell Decoder aims to identify cell types from gene expressions by explicitly modelling the multi-scale biological interactions, i.e., genes, pathways and biological processes. Specifically, Cell Decoder designs intra-scale and inter-scale graph neural network (GNN) layers to learn the complex patterns of multi-scale structures and properties through message passing^58^. Additionally, Cell Decoder incorporates AutoML techniques to automatically design the intra-scale and inter-scale GNN layers as well as the hyper-parameters, thereby enhancing the adaptivity of the model in handling different application scenarios. To further improve the interpretability, Cell Decoder adds post-hoc interpretability modules to provide explainable analyses for both features and multi-scale interactions (Supplementary Fig. 5).

#### Multi-scale Graph

Cell Decoder captures the interactions among genes, pathways and biological processes using a multi-scale graph. The gene expressions and interactions between genes are represented as an undirected graph with an adjacency matrix **A**^gn^ ∈ {0,1}^*n*×*n*^and a feature matrix **X** ∈ ℝ^*n*×*f*^, where *n* denotes the number of genes and 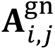 indicates whether the *i*-th gene interacts with the *j*-th gene. Cell Decoder also considers the gene-pathway relationship **A**^gn→pw^, where 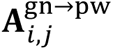 indicates whether the *i*-th gene belongs to the *j* -th pathway. Similarly, the pathway-biological process relationship is taken into account and denoted as **A**^pw→bp^. Then, Cell Decoder constructs the pathway adjacency matrix by utilising the multi-scale biological knowledge:

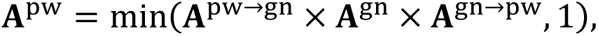

where the operator × is the matrix multiplication, and **A**^pw→gn^ is the transpose of **A**^gn→pw^. The biological process adjacency matrix is similarly constructed, i.e.,

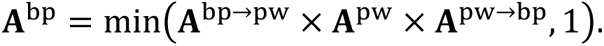

#### Multi-scale Message Passing

The objective of the multi-scale message-passing layers in Cell Decoder is to effectively aggregate information from genes, pathways, and biological processes. To incorporate multi-scale information, Cell Decoder consists of two types of GNN layers: intra-scale layers and inter-scale layers. For intra-scale layers, Cell Decoder aggregates messages from the neighborhoods within each scale and subsequently updates the node representations as follows:

**H**^gn^ ← GNN_Intra_(**H**^gn^, **A**^gn^), **H**^pw^ ← GNN_Intra_(**H**^pw^, **A**^pw^), **H**^bp^ ← GNN_Intra_(**H**^bp^, **A**^bp^), where **H** ∈ ℝ^*n*×*d*^ is the node representation, and *d* denotes the dimensionality. For inter-scale layers, Cell Decoder updates the node representations by aggregating the neighborhood messages from preceding scales, i.e.,

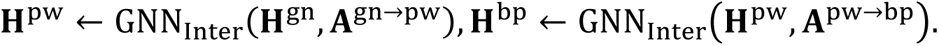

The execution of intra-scale layers and inter-scale layers follows an alternating pattern, starting from genes, then pathways, and finally biological processes. Finally, Cell Decoder utilises mean pooling to summarise the node representations into cell representations, which are subsequently employed for cell-type identification, i.e.,

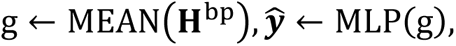

where g ∈ ℝ^1×*d*^ represents the cell representation, MLP denotes a multi-layer perceptron classifier, MEAN(⋅) is the mean pooling operator,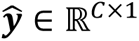 represents the predicted probability distribution of the cell types, and *C* is the number of the cell types. To learn the model parameters, Cell Decoder adopts the cross-entropy loss function:

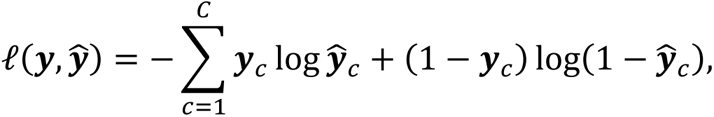

where ***y*** denotes the ground-truth cell types.

#### Automated Model Design

Given that biological mechanisms in various cells could show intricate patterns, it becomes crucial to identify optimal hyper-parameters and GNN layers for improved cell-type identification. In this regard, we employ AutoML techniques to automate the process of architecture design and hyper-parameter selection. To maintain simplicity, we adopt a classical grid search method^59^, which systematically enumerates and selects the best architectures and hyper-parameters from the entire set of possibilities. The search space of Cell Decoder encompasses intra-scale layers, inter-scale layers, hyper-parameters, and other architectural modifications, defining the feasible ranges for exploration and selection.

#### Intra-scale Layers

Our search space includes two types of intra-scale layers: Graph Attention Networks (GAT)^60^ and Graph Isomorphism Networks (GIN)^61^.

GAT employs an attention mechanism for neighborhood aggregation, and its formulation is as follows:

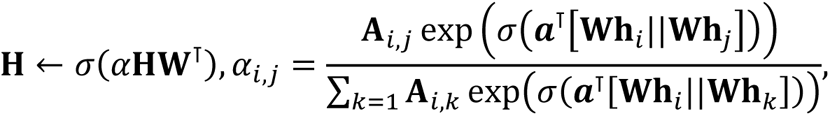

where **W** represents a learnable parameter matrix used to transform the node representations, ***a*** is a learnable vector for calculating the attention weights *α* for each edge, (⋅)^⊺^denotes the transpose operation, **h**_*i*_ denotes the feature vector of the *i* -th node, || denotes the concatenation operation, and *σ* represents the LeakyReLU activation function. Multi-head attention mechanism is also utilised to stabilise the training process and improve performance.

On the other hand, GIN, which adopts a sum aggregation, can be formulated as follows:

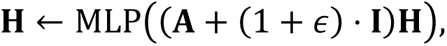

where *ϵ* is a learnable parameter, **I** denotes the identity matrix, and MLP represents a multi-layer perceptron with learnable parameters.

#### Inter-scale Layers

For inter-scale layers, our search space contains two operations: mean pooling^62^ and attention pooling^63^.

The mean pooling computes the average of the neighbour representations from the previous layers to obtain the node representation in the current layer. For instance, to obtain the representation of the *i*-th pathway in gene-pathway message-passing, the model aggregates all of its related gene representations and calculates the average as follows:

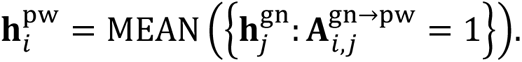

In the case of attention pooling, the calculation is similar to Eq. (7), with the difference that we first employ mean pooling to obtain the node representation used in calculating attention, and then utilise the inter-scale adjacency matrix to calculate and aggregate the messages. For example, the hidden representation for the pathways is calculated as:

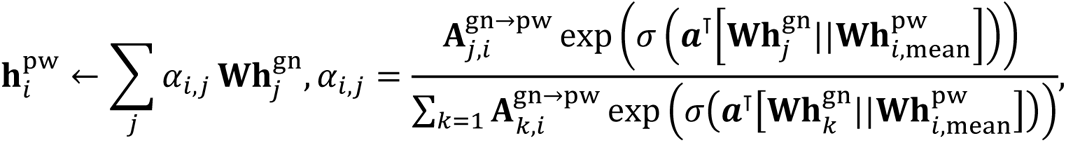

where 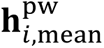 is calculated by the mean pooling.

#### Hyper-parameters

Additionally, our search space encompasses commonly used hyper-parameters, including the dimensionality of the node representation, the learning rate, and the number of samples per class utilised for training. If there exist some classes that have fewer or more training samples than *K*, we conduct over-sampling or down-sampling techniques to obtain exact *K* training samples for each class.

#### Architecture Modification

Lastly, our search space includes two architecture modification techniques to facilitate more flexible architecture design: one-hot encoding of genes and jumping connections.

For the one-hot encoding, we introduce a learnable embedding matrix for each gene node, which is concatenated with the node features as follows:

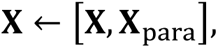

where **X**_para_ denotes the trainable embedding matrix. This enables the model to effectively capture specific information pertaining to each gene, enhancing its ability to learn gene-specific characteristics. Regarding the jumping connections, we allow the model to concatenate the raw node features with the final node representation learned by the GNN layers. The concatenated features are then fed into the classifier as follows:

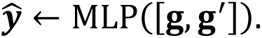

where **g**′ is obtained by flattening the raw features **X** for each graph into a vector. This strategy draws inspiration from the widely employed residual connections in deep learning^64^ and the jumping knowledge network in GNNs^65^. By incorporating jumping connections, we aim to improve the expressiveness of the model and enhance training stability.

### Multi-view Model Interpretability

In order to interpret the model, we employ three types of post-hoc analysis to assess the importance of features, edges, and cross-edges, respectively.

#### Feature Importance

We utilise two methods to measure the feature importance: gradient-based methods (referred to as Grad) and Gradient-weighted Class Activation Mapping (GradCAM)^20^. For Grad, we calculate the L2 norm of the feature gradients with respect to the loss, which quantifies the model’s sensitivity to the features. GradCAM extends this analysis by incorporating the feature maps prior to graph convolutions, taking into account the influence of features on the classifiers.

#### Edge Importance

Similar to GNNExplainer^66^, we utilise learnable edge masks to evaluate the importance of edges between genes in the model. These masks are learned by optimising the following objective:

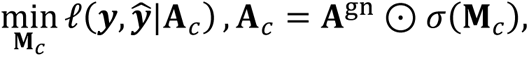

where **M**_*c*_ denotes the mask for class *c*, *σ*(⋅) is the sigmoid activation function, ⊙ denotes the element-wise matrix product, and 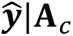 is the cell type prediction when using the masked adjacency matrix **A**_*c*_. The rationale behind this approach is that if certain masked edges have minimal impact on cell type prediction, their importance to the model is relatively low, and vice versa.

### Cross-edge Importance

When Cell Decoder utilises attention pooling in the inter-scale layers, we leverage the attention weights in pooling functions to directly assess the importance of cross-edges. If the attention weight *α*_*i*,*j*_ is large, it indicates that the edge connecting the *i*-th node in the preceding layer and the *j*-th node in the current layer holds significance, since a considerable weight is assigned to this edge during the aggregation of neighbourhood messages.

### Robustness Evaluation

We evaluate the robustness of our model through three perturbation strategies, which involve perturbation on features, nodes and structures. We focus on the robustness of model ^67^, where all perturbation occur during test time by manipulating the input data.

**Feature** perturbation. To perturb a feature *x*, we denote its mean and standard deviation as *μ* and *σ*, respectively. The feature is modified as *x* ← max(*x* + *λ* ⋅ *ϵ*, 0), where the noise *ϵ* is sampled from a Gaussian distribution 𝒩(*μ*, *σ*), and *λ* ∈ ℝ is a hyper-parameter that controls the perturbation rate.

**Node** perturbation. In this perturbation, we randomly remove nodes and denote *p* ∈ [0,100] as a hyper-parameter to control the perturbation rate. We randomly select *p*% of the gene nodes and remove all their edges from the graph, except for the self-loops.

**Structure** perturbation. In this perturbation, we randomly remove edges and denote *p* ∈ [0,100] as a hyper-parameter to control the perturbation rate. We randomly select *p*% of the edges from the adjacency matrices **A**^gn^, **A**^gn→pw^, **A**^pw→bp^, and remove these edges from the graph.

### Datasets and Pre-processing

All datasets utilised in this study are obtained from public data repositories (Supplementary Data 1). All datasets used for cell-type identification benchmarking were computed with 3000 highly variable genes (HVGs) using scanpy^68^.

### Baselines for cell identification and data integration

All methods used for cell-type identification comparison were trained on the same training set with default recommended parameters. In the data integration task, integrations were evaluated with methods implemented in scIB using default recommended parameters.

### Metrics for cell identification evaluation and integration

We use accuracy and Macro F1 to evaluate the performance and robustness of models in predicting cell types. For the data integration task, we employ average silhouette width (ASW), Batch ASW, kBET, Graph Connectivity and Graph iLISI to assess the model’s batch correction capability. Additionally, we utilise normalised mutual information (NMI), adjusted Rand index (ARI), Cell Type ASW, and Isolated Label F1 to evaluate the model’s ability to preserve biological differences using scIB. The overall score was computed as the average of all scores.

### Statistics and Reproducibility

The details for data pre-processing of datasets are provided in the section ‘Datasets and pre-processing’. Unless otherwise specified in the respective section or figure legends, all data were included in the training and analysis. The architecture and hyperparameters for model training are provided in Supplementary Data 5.

## Data Availability

Datasets used for cell identification benchmarking are available from GEO (GSE171555, GSE134355, GSE136103, GSE115469, GSE145927, GSE81608, GSE84133, GSE132188, GSE252225) and ArrayExpress (E-MTAB-5061). The human immune cell datasets used for integration analysis are downloaded from 10X Genomics website (https://support.10xgenomics.com/single-cell-gene-expression/datasets/3.0.0/pbmc_10k_v3) and GEO (GSE120221, GSE107727, GSE115189, GSE128066, GSE94820). The mouse embryo datasets are from GEO (GSE109071, GSE100597) and ArrayExpress (E-MTAB-6967). More detailed information can be found in the supplementary data.

## Acknowledgements

We would like to acknowledge Dr. Fan Zhou (Tsinghua University) for his insightful comments. J.Z., L.X., F.H. and C.C. are financially supported by the National Key Research and Development Program of China (2021YFA1301603 and 2020YFE0202200) and the National Natural Science Foundation of China (32088101). Z.Z. and W.Z. are supported by the National Key Research and Development Program of China (2020AAA0106300) and the National Natural Science Foundation of China (62250008). Z.Z. is also supported by the National Natural Science Foundation of China (62206149).

## Author Contributions

J.Z. and Zeyang.Z. conceived the project and designed the architecture of Cell Decoder. C.C. and Ziwei.Z. co-supervised the project. J.Z., Y.X. and B.X. performed data analysis. X.D., R.R., X.W. and Y.L. collected the public datasets. F.H. and W.Z. coordinated the study. L.X. and P.Z. helped to revise the manuscript. All authors contributed to the writing of the manuscript.

