## Supplementary Information for "Decoding cell identity with multi-scale explainable deep learning"

**of**

### Supplementary Figures

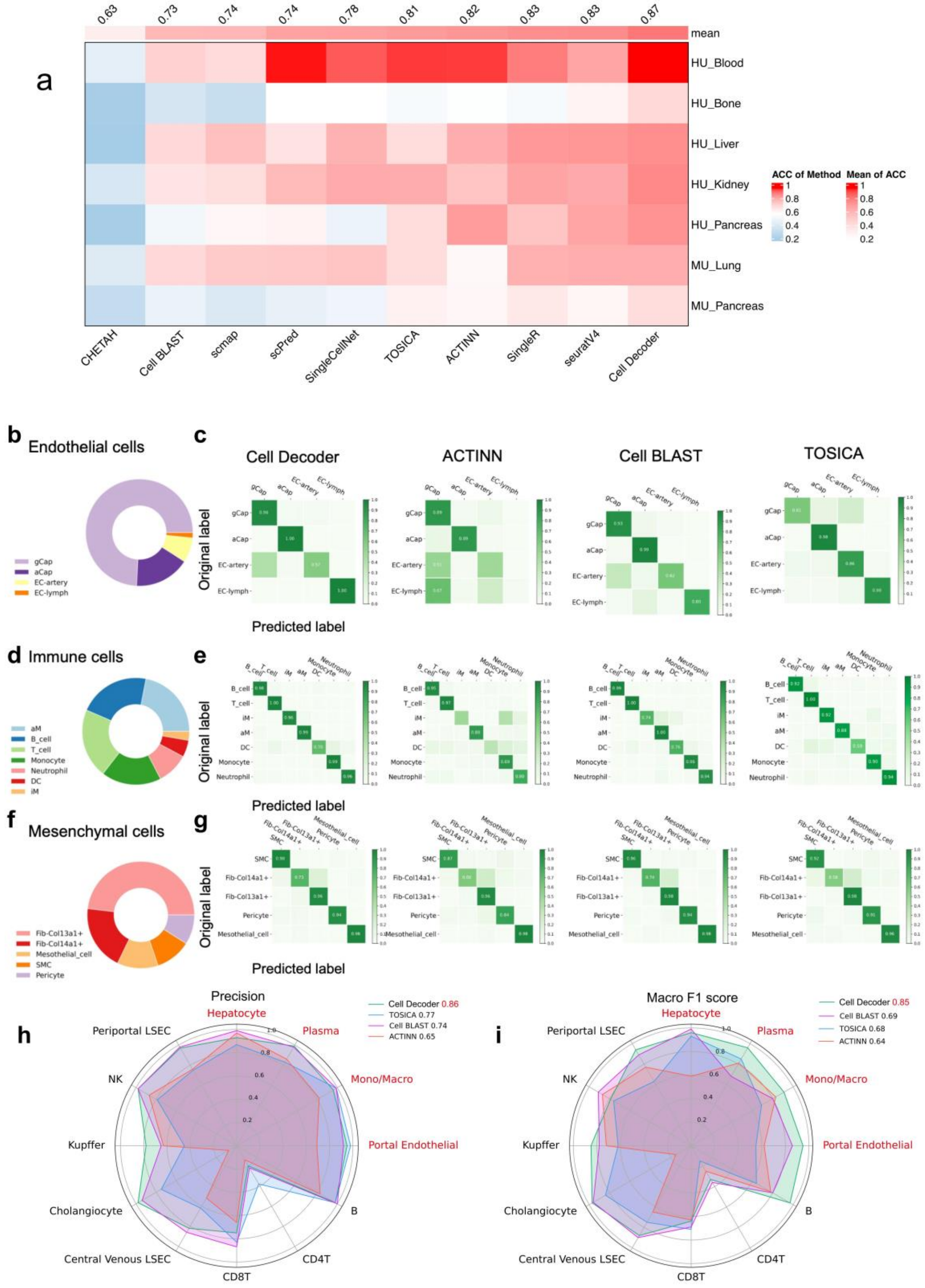


**Supplementary Fig. 1. Benchmarks of Cell Decoder for cell identification. a** Accuracy for different cell identification methods on human and mouse datasets. Columns are sorted by the mean accuracy (ACC) of each method on all datasets (top). **b-g** The prediction performance of Cell Decoder on imbalanced cell types is shown in the confusion matrix. Rows represent the original labels, and columns represent the predicted labels. Specifically, **b-c** for endothelial cells, **d-e** for immune cells, and **f-g** for mesenchymal cells. **h-i** Precision and Macro F1 score of each cell type (indicated in red if its proportion in the query is greater than that in the reference), and the average Precision and Macro F1 scores for each method are labelled on the upper right corner.

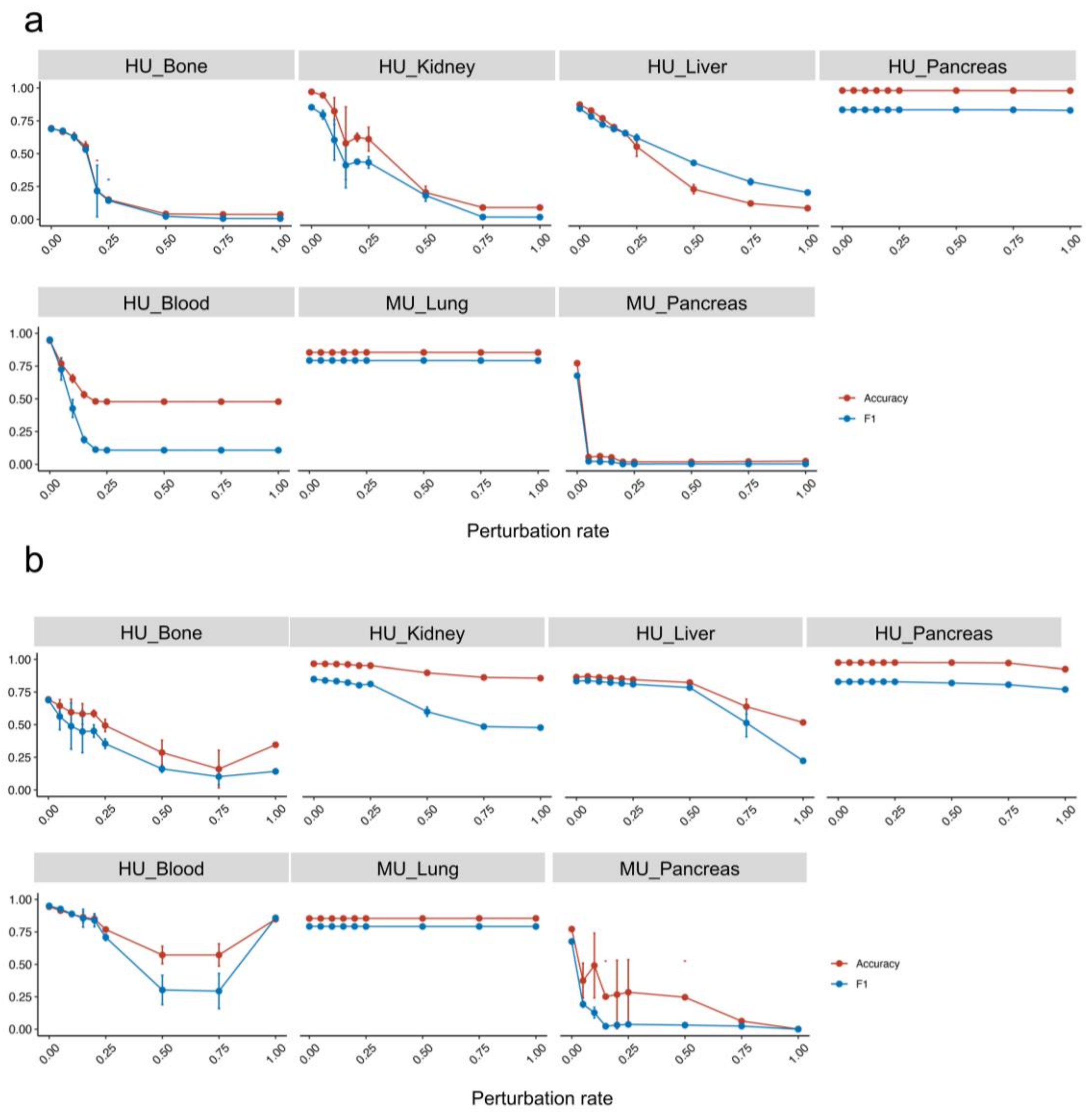


**Supplementary Fig. 2. Biological knowledge ablation experiments for Cell Decoder**. **a** Random node ablation experiments were conducted by randomly removing a certain proportion of nodes from the network. The x-axis represents the perturbation ratio, while the y-axis represents the post-perturbation prediction accuracy (red line) and Macro F1 score (blue line). **b** Edge ablation experiments were conducted by randomly removing existing edges in the network at a certain proportion. The x-axis represents the perturbation ratio, while the y-axis represents the post-perturbation prediction accuracy (red line) and Macro F1 score (blue line).

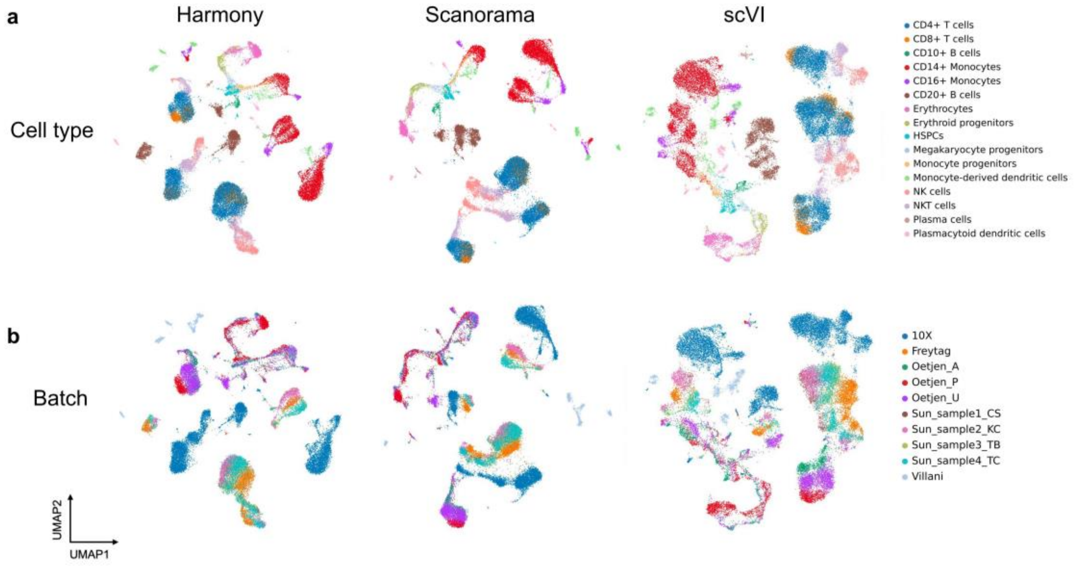


**Supplementary Fig. 3. UMAP plots for human immune datasets.** Visualisation of the Harmony, Scanorama and scVI on the immune cell integration without batch labels coloured by cell type (**a**) and batch annotation (**b**).

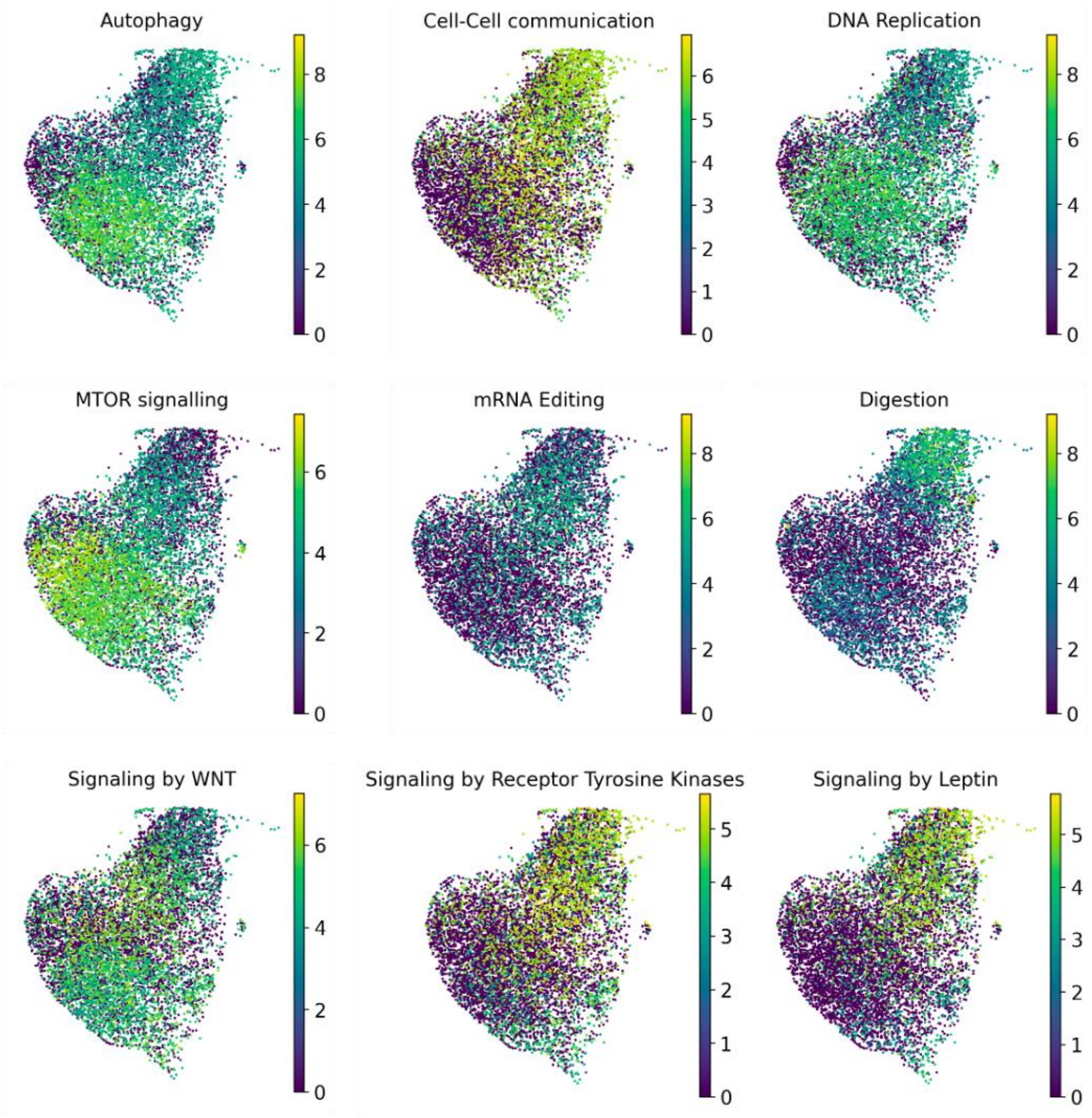


**Supplementary Fig. 4. The distribution of the representative pathways with top Grad-CAM scores across the three cell states of the HU_Bone dataset.** This figure is coloured by the Grad-CAM score of a pathway.

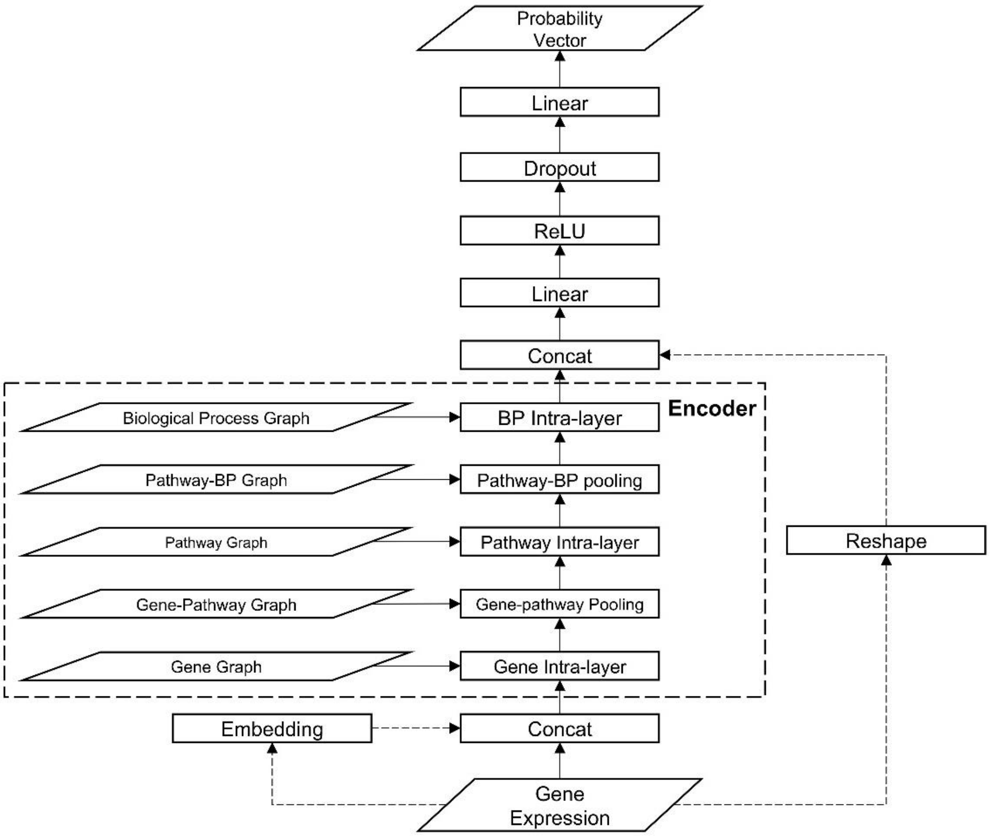


**Supplementary Fig. 5. Architecture of Cell Decoder in detail.**
The gene expression profile of one cell initially undergoes encoding or optionally gets concatenated with an embedding vector before being processed through the encoder. This encoder phase includes intra-layer operations and pooling. Subsequently, the output from the encoder serves as input for a linear classifier. Dotted lines indicate optional architecture in automated machine learning. Please refer to the methods section for further elaboration.
